# Chimpanzee fibroblasts exhibit greater adherence and migratory phenotypes than human fibroblasts

**DOI:** 10.1101/838755

**Authors:** Trisha M. Zintel, Delaney Ducey, Courtney C. Babbitt

## Abstract

**Background and objectives:** Previous work has identified that gene expression differences in cell adhesion pathways exist between humans and chimpanzees. Here, we used a comparative cell biology approach to assay interspecies differences in cell adhesion phenotypes in order to better understand the basic biological differences between species’ epithelial cells that may underly the organism-level differences we see in wound healing and cancer.

**Methodology:** We used skin fibroblast cell lines from humans and chimpanzees to assay cell adhesion and migration. We then utilized published RNA-Seq data from the same cell lines exposed to a cancer / wound-healing mimic to determine what gene expression changes may be corresponding to altered cellular adhesion dynamics between species.

**Results:** The functional adhesion and migration assays revealed that chimpanzee fibroblasts adhered sooner and remained adherent for significantly longer and move into a “wound” at faster rate than human fibroblasts. The gene expression data suggest that the enhanced adhesive properties of chimpanzee fibroblasts may be due to chimpanzee fibroblasts exhibiting significantly higher expression of cell and focal adhesion molecule genes than human cells, both during a wound healing assay and at rest.

**Conclusions and implications:** Chimpanzee fibroblasts exhibit stronger adhesion and greater cell migration than human fibroblasts. This may be due to divergent gene expression of focal adhesion and cell adhesion molecules, such as integrins, laminins, and cadherins, as well as ECM proteins like collagens. This is one of few studies demonstrating that these divergences in gene expression between closely related species can manifest in fundamental differences in cell biology. Our results provide better insight into species-specific cell biology phenotypes and how they may influence more complex traits, such as cancer metastasis and wound healing.

## INTRODUCTION

Although humans and chimpanzees shared a common ancestor relatively recently (∼ 5 million years ago), they have largely identical amino acid and protein structure and greater than 98% DNA sequence identity [1, 2]. Despite this high nucleotide conservation, there are many well-documented phenotypic discrepancies related to cognition, behavior, and anatomy, including some intriguing differences that may be biomedically relevant for understanding human-specific disease [1, 3]. Determining specific genetic influences on interspecies disparities in disease etiology is important to understand human-specific diseases [4, 5]. In order to better understand these organism-level differences in complex phenotypes, and their evolutionary basis, we can investigate interspecies differences in basic molecular and cellular biological processes.

One mechanism by which interspecies differences in phenotype may arise from highly similar genomes is through changes in non-coding sequences that influence the expression the same protein-coding genes [2]. Significant changes in gene expression between humans and chimpanzees have been studied previously [6-14]; however, studies of the cellular phenotypic impacts of altered gene expression are lacking. One way to start to make those connections is to use available cell lines from multiple species. Fibroblast cells are important for homeostasis and critical for wound healing and often play an integral role in inflammation during cancer progression [15, 16]. A previous publication found that chimpanzee skin fibroblasts have significantly more focal adhesions per cell than human fibroblasts [17]. Additionally, human and chimpanzee fibroblasts exhibit significant differences in gene expression of adhesion pathways both in fibroblasts grown under normal conditions [11, 18] as well as those exposed to a serum challenge that mimics both wound healing and a cancer response [18]. However, there are no known studies investigating if these differences influence cellular adhesion and migratory phenotypes.

Intercellular signaling facilitated by cell adhesion is vital for many biological processes, such as embryonic development as well as disease states (e.g. metastasis of cancerous cells) and consequently represents a potential mechanism by which functional differences among primates may have evolved. Healing of a wound *in vivo* requires migration of neighboring healthy cells into the wound [19] which is facilitated by disassembly and reassembly of focal adhesion complexes to the leading edge of migrating cells [20-23]. Cell adhesion and migration pathways are important in a number of complex traits, including development and differentiation [23-26] as well as cancer metastasis [27-30]. Focal adhesions (FAs) are large protein complexes that facilitate adhesion of cells to the underlying extracellular matrix (ECM) as well as cell motility by the disassembly and reassembly of these complexes in difference locations along the cell’s adherent surface [20-23, 31]. A variety of protein types comprise FAs, including extracellular protein complexes (e.g. zyxin) that interact extracellular matrix proteins (e.g. collagen, laminin) to facilitate adherence as well as transmembrane proteins that function not only in adherence but also in bidirectional signaling across the membrane (e.g. vinculin, integrin) [20].

In order to determine if previous findings of interspecies differences in adhesion complexes and gene expression translate into fundamental differences in cellular phenotypes, we used established cell culture methods to assay adhesion and migration differences between human and chimpanzee primary skin fibroblasts *in vitro*. We hypothesized that there may be significant differences in cellular adhesion and migration phenotypes between human and chimpanzee fibroblasts that correlate with the previously observed differences in focal adhesion gene expression. We investigated adhesive properties using two different cell adhesion assays, that measured timing and strength of fibroblast adherence and migration phenotypes by comparing migration patterns, speed, and distance of cells over time in two conditions. We also used a classic scratch test that mimics a wound and the subsequent healing process *in vitro* [32, 33]. Finally, in order to examine underlying gene expression differences in adhesion and migration signaling pathways between species at the cellular level, we incorporated previously published data on differential gene expression between human and chimpanzee fibroblasts during an assay that mimics both wound healing and cancer progression [18].

## METHODOLOGY

### Samples

Primary fibroblasts from two individuals per species were obtained from Coriell Biorepository (Camden, NJ) (human lines AG10803 and AG09605, chimpanzee lines S006007 and S008956). Cells were cultured with minimum essential media (MEM) supplemented with 10% fetal bovine serum (FBS), 1% L-glutamine, & .1% antibiotic at 37C and 5% CO_2_, according to Coriell protocols. Cells were sex-, age-, and body-region-matched.

### Cell adhesion assay

Fibroblasts at 80-90% confluency in 6-well plates were dyed with a non-toxic cytoplasmic dye for ten minutes at a concentration of 1 μM (catalog #C7025, Invitrogen). A live cell nuclear stain was then added for twenty minutes, according to manufacturer’s instructions (#R37605, Invitrogen). Fibroblasts were subsequently passaged using .5% trypsin and seeded as evenly as possible into eight individual wells of a 12-well plate. At four timepoints (.5, 1, 1.5, 2 hours), wells were rinsed to remove non-adherent cells and fluorescent images were taken. To maximize accurate counts of cells/well, each well was divided into two quadrants, and for each timepoint, one photo per quadrant was taken. Cell counts were obtained using FIJI Is Just ImageJ (FIJI) [34] to quantify the fluorescently dyed cells. We calculated a mean of the two counts per timepoint. Experiments were conducted simultaneously in triplicate for all four cell lines. DNA concentration was used to control for differences in cell number by dividing cell counts at each time point by the DNA concentration an equal aliquot from time zero.

### Flow challenge to assay cell adhesion and migration

Fibroblasts of each species were assayed for differences in cell adherence and migration using a microfluidic device equipped with a time-capture, environment-controlled microscope system, developed in Shelly Peyton’s laboratory (University of Massachusetts Amherst). Fibroblasts at 80% confluency were trypsinized and 1,000-150,000 cells were seeded into the flow cell device to monitor cell adhesion and migration over time (referred to further as adhesion-in-flow experiments). The flow cell device uses two reciprocating syringe pumps connected via tubing to a narrow chamber over which cells suspended in media were pumped into at a rate of 50v µL/minute for 6 hours. Photographs of the fibroblasts adhering and migrating in the flow cell were taken every 15 minutes over the 6-hour time course. Individual fibroblasts were identified and assessed over the 6-hour observation period and were traced using FIJI consecutively across this time period to obtain measurements of initial fibroblast adherent surface area and subsequent cell spread via centroid X and Y coordinates over time [34]. These data were used to plot migration paths of individual fibroblasts per species (SI Figure 1A). Distance traveled of individual fibroblasts was calculated with a standard distance equation (D = [(x_2_-x_1_)_2_ + (y_2_-y_1_)_2_]) [35]. Speed was calculated in microns/minute using total distance traveled divided by time spent adherent for each individual cell [36]. Adherence duration was calculated by averaging total lengths of time individual fibroblasts spent adherent. Timing of fibroblast adherence was determined by comparing the proportion of fibroblasts adherent at a particular time point (i.e. .5 hours, 1 hour) to that of the total number of fibroblasts that adhered over the entire time course. However, due to significant cell death, sufficient cell numbers (n = 25-30, per experiment) were obtained only for a single experiment per cell line (two individuals per species) and we could not obtain the planned technical replicates per cell line we had attempted. However, per species, we analyzed a minimum of 80 cells. Significant cell death likely occurred due to the required changes from typical mammalian cell culture needed for this assay, which included a transition from CO_2_-independent media to CO_2_-dependent media in addition to consistent sheer force on these primary, un-immortalized cells due to the reciprocating flow across the surface for the duration of the experiment.

### Wound healing assay

In order to assess interspecies differences in cell migration, we conducted a wound healing assay as previously described [32, 33]. Briefly, 70-90% confluent fibroblasts were dyed with a non-toxic cytoplasmic dye as previously described and scratched (“wounded”) using a p200 pipette tip. Each well was then rinsed and media was replaced to remove cell debris prior to taking fluorescent images (time 0). The same region of the well was subsequently imaged at 24, 48, and 72 hours. We used CellProfiler (cellprofiler.org) to quantify the number of cells that migrated into the wound and the area covered by cells over time [37]. Experiments were conducted in triplicate for all four cell lines.

### Identifying candidates for altered cell adhesion and migration from a study of differential expression of genes during a wound healing response

In order to determine gene expression differences that may correspond to the interspecies cellular differences in adhesion in our experiments, we used data produced by Pizzollo et al. [18], a published study from our laboratory, that determined genes exhibiting significant different expression (DE) (q-value < 0.05) between human and chimpanzee fibroblasts undergoing a wound healing response. This study investigated global open chromatin and transcription (via DNAse-Seq and RNA-Seq, respectively) in the same cell lines investigated here. Pizzollo et al. [18] found genes exhibiting significant DE between species’ fibroblasts at rest (unchallenged) and those undergoing a similar wound-healing response to the one used in our experiments. For our analysis, we obtained these lists of significantly differentially expressed (DE) genes from [18] and narrowed them down to those that are members of the Kyoto Encyclopedia of Genes and Genomes (KEGG) pathways [38] for “focal adhesion” and “cell adhesion molecules” (SI Table 1). We then compared whether they were DE between species fibroblasts at rest, during the wound healing response, or both to determine putative genetic differences underling the cellular phenotype differences in adhesion and migration observed in our experiments.

## RESULTS

### There are phenotypic differences in cell adhesion between human and chimpanzee fibroblasts

In order to investigate if cellular phenotypes related to adhesion differ in manner consistent with previously findings of significant differences in focal adhesions between human and chimpanzee fibroblasts [17], we wanted to determine if there were differences in timing of fibroblast adherence to a surface. In order to determine if there was a difference between species in incidence of adhesion over time, we conducted multiple assays of fibroblast adhesion and compared across species. Initially, we conducted a straightforward investigation of cell adherence by seeding trypsonized fibroblasts into multiple wells and then removing excess, suspended cells at sequential timepoints (.5, 1, 1.5, and 2 hours post-seeding) to count the number of cells that had already adhered (Figure 1A). We expected that if chimpanzee cells adhered more strongly than human cells, we would see more chimpanzee cells adhere during earlier timepoints. We observed greater numbers of chimpanzee cells adhering than human fibroblasts at all timepoints (Figure 1A). There was a statistically significant difference between species (two-way ANOVA F=13.59, p = 0.0142) but not time (two-way ANOVA F=.62, p = 0.4665) (Figure 1A).

**Figure 1.**
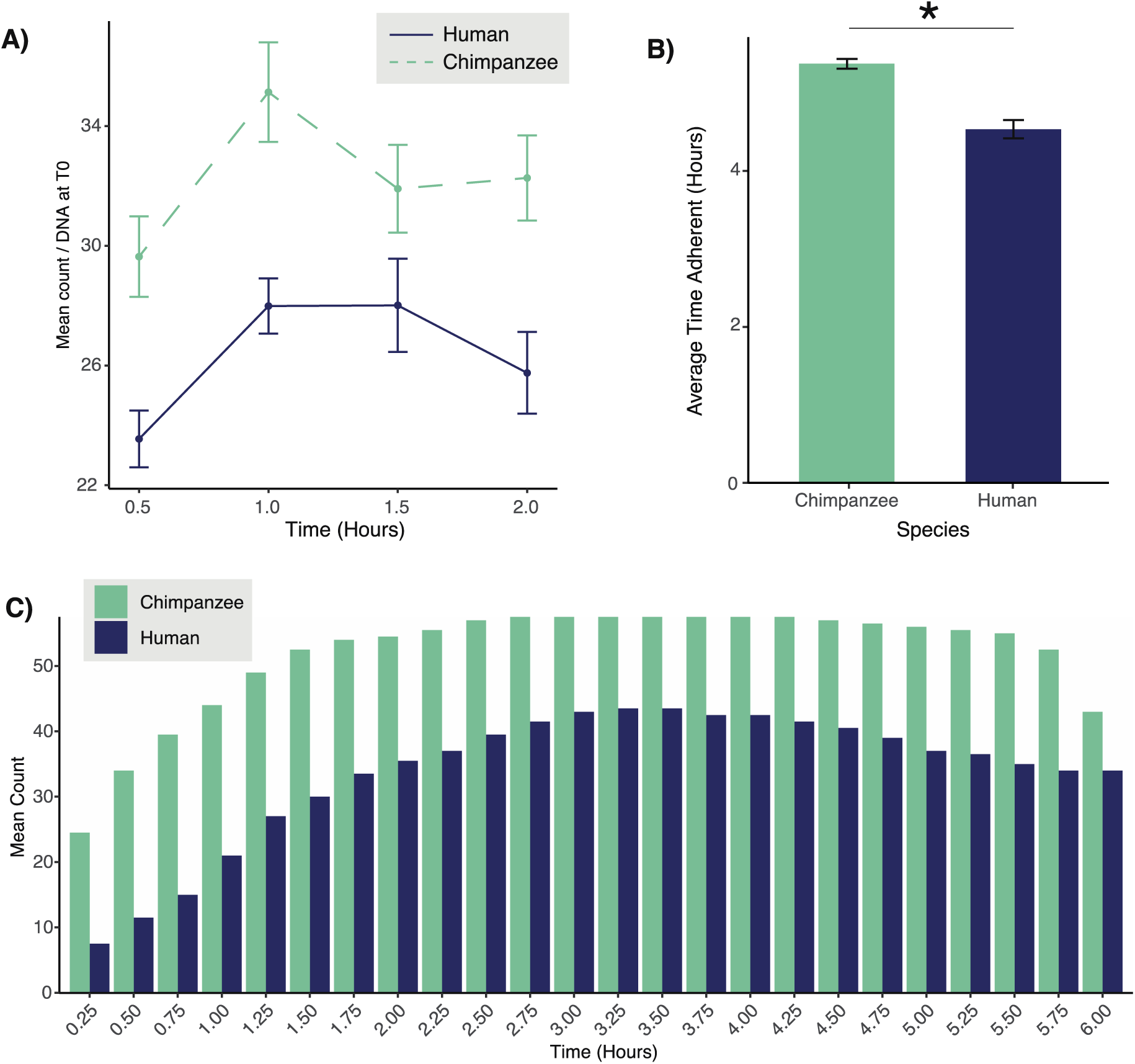
Adhesion differences between human and chimpanzee fibroblasts. Fibroblasts from two individuals per species were assayed for adherence *in static* (A) and *in flow* (B, C). A) Adhesion-in-static assay. Mean count per timepoint normalized for differences in cell number by dividing raw cell counts at each time point by the DNA concentration from time zero. B) Comparison of the duration of fibroblast adherence *in flow* (per species) (p < .001; Welch 2-sample T-test). C) Comparison between species of the timing of fibroblast adherence over the 6-hour time course.

We next used a more complex assay for cell adhesion using a flow cell device that continuously flowed media with suspended fibroblasts back and forth over a plastic surface, allowing us to monitor adhesion via photographs over a six hour time period (further referred to as the ‘adhesion-in-flow’ assay). One benefit to this approach was that that allowed us to better control cell number across experiments (100,000 per experiment for human cells, 150,000 for chimpanzee cells). Individual fibroblasts were identified in these sets of images and traced over time to obtain counts of adherent cells. We calculated percentages of observed adherent cells at each time point as well as the length of time cells of each species remained adherent. When we calculated the mean total length of time fibroblasts were adherent over the six hour time course, chimpanzee fibroblasts were adherent for significantly longer (5.4 hours) than human fibroblasts (4.56 hours, p < .001; Welch 2-sample T-test; Figure 1B) and adhered in greater numbers within the first hour than human cells (Figure 1C). These results demonstrate chimpanzee fibroblasts adhering faster than human fibroblasts and may mean chimpanzee cells exhibit firmer cell adhesion than human fibroblasts.

### Chimpanzee fibroblasts do not appear to inherently migrate faster or farther than human fibroblasts but do exhibit faster *in vitro* wound healing

The increased number of focal adhesions in chimpanzee fibroblasts may also confer differences in cell motility due to dynamic nature of disassembly and reassembly of focal adhesion complexes during cell migration [20-23]. We investigated if there were inherent differences in cell migration between species. Individual fibroblasts from the adhesion-in-flow experiments were traced using FIJI software throughout the full six-hour experiment to obtain area measurements and X & Y coordinates. In order to determine if there were differences in migration patterns between fibroblasts, we used the centroid X and Y coordinates obtained from the six-hour set of photographs to construct a plot of migration patterns between species (SI Figure 1A). There appears to be little interspecies difference in migration patterns (SI Figure 1A), though, qualitatively, there does seem to be more movement along the Y-axis for human cells than chimpanzee cells. Using these coordinates, we calculated distance traveled and speed of migration for each individual fibroblast over time. There was not a significant difference between species in mean distance migrated (SI Figure 1B) or speed of migration (microns/hour, SI Figure 1C). There was also no difference in cell spread (area/time, data not shown).

Given this fundamental biological link between focal adhesions and cell motility as well as the evidence for greater adhesive properties of chimpanzee cells, we further assessed cell migration by conducting wound healing experiments, which allow us to test a more biologically relevant scenario of cell migration (SI Figure 1A). Wound healing assays, also known as ‘scratch tests’ are a straightforward way to investigate cell migration *in vitro*, and involve making a scratch in a monolayer of cultured cells and monitoring the scratch over time as cells respond to the gap [32, 33]. We found a statistically significantly increase in number of chimpanzee fibroblasts that migrated into the wound (two-way ANOVA F=30.36, p = 1.66e-06; Figure 2B). There was also a statistically significant difference in cell count for both species in original scratch over time (two-way ANOVA F=62.02, p = 5.23e-10; Figure 2B). This resulted in chimpanzee fibroblasts covering a greater percentage of the initial wounded area than human fibroblasts during the first 24 hours post-wounding (Figure 2C). However, while there was a statistically significant difference in area covered over time (two-way ANOVA F=14.20, p = 0.000475), there was not between species (two-way ANOVA F=1.18, p = 0.283221). These results demonstrate that larger numbers of chimpanzee fibroblasts migrate into an *in vitro* wound than human fibroblasts do.

**Figure 2.**
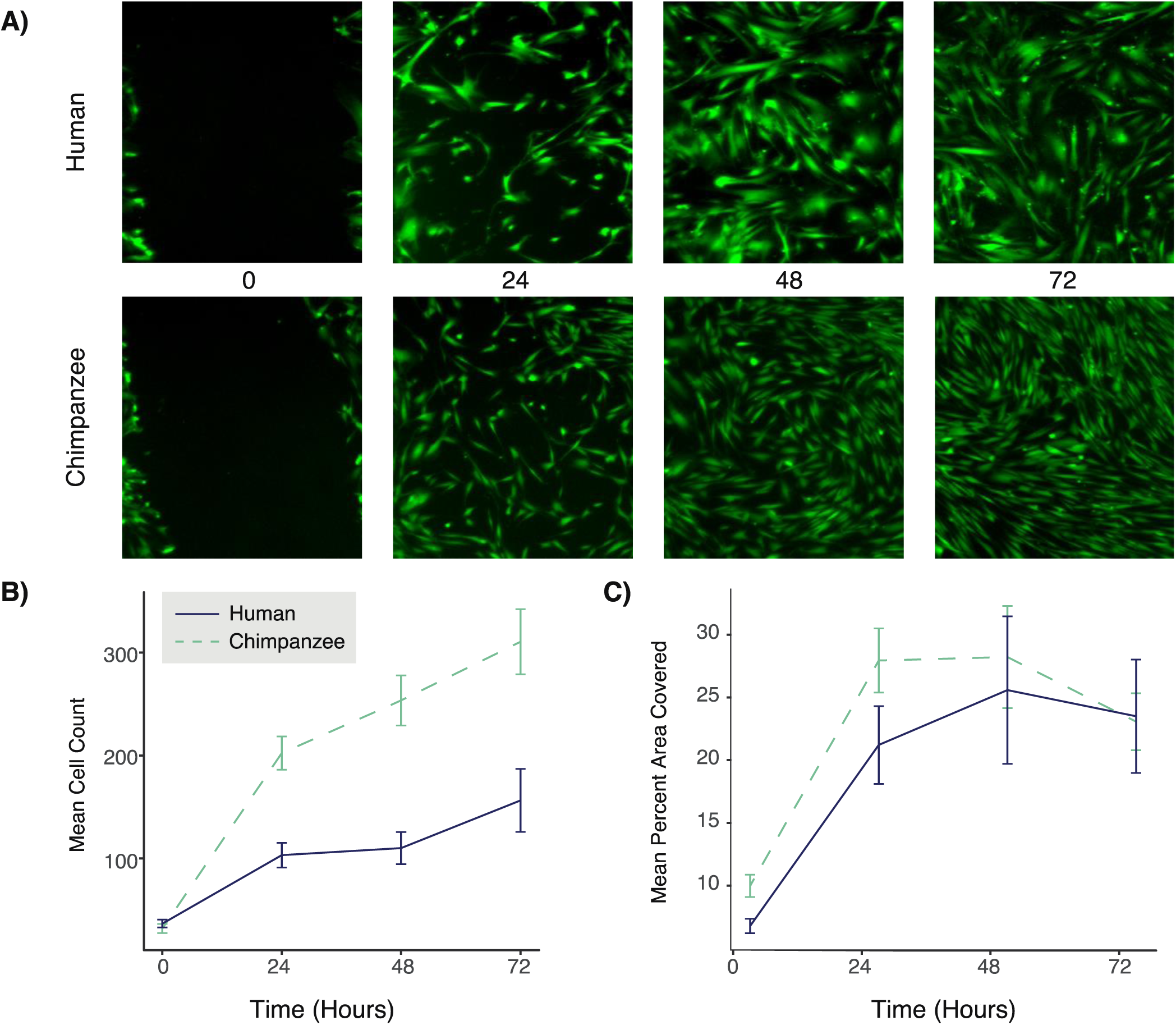
Interspecies differences in wound healing. A) Representative images of human and chimpanzee fibroblasts undergoing a wound healing response after initial scratch (0), and 24, 48, and 72 hours post-scratch. CellProfiler was used to quantify the number of cells that moved in to fill in the scratch (B) and the change in area covered by cells (C) at 0, 24, 48, and 72 hours post-scratch.

### Human and chimpanzee fibroblasts increase expression of distinct FA and CAM genes normally and during wound healing

In order to see how gene expression diverges during the wound healing response between species, we harnessed previously published data from our lab that compared gene expression profiles via RNA-Seq of human and chimpanzee fibroblasts during a serum challenge that mimics both wound healing and metastasis, using many of the same cell lines as those investigated here [18]. In this study, while human and chimpanzee fibroblasts both exhibited gene expression profiles characteristic of wound healing throughout the serum challenge, there were a number of genes that were significantly differentially expressed (DE) between species in the control, unchallenged samples (pre-challenge) as well as during the wound healing response (SI Table 1) [18]. In order to determine what interspecies differential expression might explain differences in adhesive properties between normal primary fibroblasts, and may be coinciding with our observed increase in cell migration during the wound healing assay, we identified genes involved in the KEGG “focal adhesion” (FA) and “cell adhesion molecules (CAMs)” pathways from the lists of genes Pizzollo et al. (2018) identified as significantly differentially expressed (DE) between species’ in the normal, unchallenged fibroblasts and during the serum response (12 and 24 hours post-serum reintroduction) [18]. Cell adhesion and migration is modulated by interplay between cell adhesion molecules and focal adhesions [39], and understanding the changes in their expression during a wound healing response can provide insight into the genetic influences on these cellular level phenotypes. There were 27 FA and 31 CAM genes DE between species at any timepoint (SI Table 1). We focused on those that belonged to functionally distinct adhesion related gene families in order to determine if there were interspecies differences expression in these gene families in unchallenged or serum-challenged fibroblasts. Several cadherin genes were DE between species at all timepoints, with CDH2 and CDH4 more highly expressed in humans and CDH1 and CDH3 more highly expressed in chimpanzees (SI Table 1,2). Chimpanzee fibroblasts exhibited significantly greater expression of the collagen gene COL4A6 at all time points and COL9A3 during the serum challenge while human fibroblasts only significantly increased expression of the collagen genes COL6A1 and COL6A3 during the serum challenge (SI Table 1,2). Chimpanzee fibroblasts had significantly higher expression of integrins at all time points (ITGA7 and ITGA9, SI Table 1,2) while human fibroblasts did not. Human fibroblasts increased LAMA4 expression during the serum challenge while chimpanzee fibroblasts always exhibited higher expression of LAMC3 and increased LAMA3 and LAMC2 when unchallenged (SI Table 1,2). Chimpanzees exhibited greater expression of the syndecan gene SDC2 at all time points, but unchallenged human cells upregulated SDC3 and while unchallenged chimpanzee cells upregulated SDC4 (SI Table 1,2). Interestingly, chimpanzees exhibited higher expression of a number of these genes at all timepoints, regardless of challenge-state (Table 1). This highlights that humans and chimpanzees are upregulating distinct genes from of FA/CAM protein families that may contribute to interspecies differences in adhesion and migration cellular phenotypes.

**Table 1.** FA and CAM genes DE between species during a serum challenge. Comparison of DE patterns of major adhesion gene family genes during serum challenge or at all time points. Asterisks indicate membership in KEGG pathway: * Cell Adhesion Molecules (CAM), ** Focal Adhesion (FA), *** both.

| DE Time | Higher in Human | Higher in Chimpanzee |
| --- | --- | --- |
| During serum challenge only | COL6A1**, COL6A3**, LAM4** | COL9A3** |
| At all time points (before & during) | CDH2*, CDH4* | CDH2*, CDH3*, COL4A6**, COL9A3**, ITGA7***, ITGA9***, LAMC3**, SDC28 |

## DISCUSSION

Our results demonstrate that chimpanzee fibroblasts adhere faster using multiple cell adhesion assays (*in static* and *in flow*) (Figures 1 & 3). We consistently observed the most stark interspecies differences at earlier time points: within the first hour for *in static* experiments (Figure 1A), the first two hours (in terms of cell count) during the adhesion-in-flow experiments (Figure 1C) likely contributing to the longer overall time adherent for chimpanzee cells (Figure 1B), as well as in the first 24 hours of the wound healing assay (Figure 2B, C). Significance in interspecies difference in cell count dropped at later time points: post-3 hours for adhesion-in-flow experiments (Figure 1C) or beyond 24 hours in the wound healing assay (Figure 2C) or became unreliable to assess in *in static* adhesion assay (Figure 1A, decrease in cell number at later times). These results in light of previous work demonstrating greater numbers of focal adhesions in chimpanzee fibroblasts [17] may suggest that biological processes dependent on rapid adherence of cells surface may differ significantly between human and chimpanzees, but those relying on more gradual adherence or cell migration may ultimately end up with similar phenotypes across species.

Given that stronger cell adhesion is associated slower cell migration [39], it is at first potentially paradoxical that we appear to observe stronger adhesion in chimpanzee fibroblasts during the early timepoints of our adhesion experiments (Fig. 1), no difference in overall cell speed or distance between species fibroblasts (Fig. 2) but apparently faster cell migration of chimpanzee fibroblasts in our wound healing assay (Fig. 3B). The biological context of these different assays likely plays a role. Our *adhesion-in-flow* experiments were stressful for these primary cells regardless of species, exemplified by high cell death (see Methods). These also assess cell migration in different contexts, one with initial cell adherence followed by migration in a challenging, flowing environment (*adhesion-in-flow*) while the wound healing scratch test assesses migration of already adherent cells into newly unoccupied space. Given that scratch/wound healing assays are a commonly used, classic assessment of cell migration, we feel these results are robust.

It appears that human fibroblasts may be larger than chimpanzee fibroblasts during our *in vitro* wound healing assay (Figure 2A). This may be responsible for the lack of significant difference between species in area covered by species in a two-way ANOVA over all times (Figure 2C). However, there is significant difference in percent of area covered at 24 hours post-scratch (Figure 2C). Our *in vitro* assessment of wound healing is not sufficient to ultimately determine if interspecies differences in cell size contribute more strongly to the anecdotal evidence of faster wound healing than number of cells migrating into wound. However, the previous work that determined that chimpanzee fibroblasts have greater numbers of focal adhesions per cell than humans controlled for differences in cell size [17], and so this does not ultimately negate these results. However, though it is not within the scope of the current study, an interesting future direction could be if the there is a larger trend toward larger cells in humans and if so, if there is any corresponding biological advantage in processes such as wound healing or cell motility.

Harnessing RNA-Seq data of the same cells lines during a similar wound-healing and cancer-like *in vitro* assay [18] allowed us to examine gene expression changes coinciding with interspecies differences in adhesion and migration during a wound healing response. We were unable to use these data to compare gene expression differences in vinculin to corroborate the finding by Advani et al. [17] of interspecies differences in vinculin staining because it was not one of the genes included in Pizzollo et al. [18]. However, it seems that interspecies differences in adhesion and migration may largely be driven by the constitutively higher expression of other CAM and FA genes in chimpanzee fibroblasts, at rest and during a wound healing response (Table 1). Cell adhesion and migration is modulated by interplay between cell adhesion molecules and focal adhesions [39] and concerted increases in expression of both likely is responsible for important interspecies differences in adhesion and cell motility. Integrins are a family of heterodimeric proteins transmembrane proteins with a critical roles in focal adhesion complexes as bidirectional signaling between the interior of the cell and the ECM, and vice versa [20, 21, 40]. It is interesting that chimpanzee fibroblasts but not human fibroblasts exhibited greater expression of integrin proteins regardless of serum status (Table 1, SI Table 2) [18]. Given that integrins are directly involved in communicating signals bidirectionally across the plasma membrane, this may indicate that chimpanzee fibroblasts have enhanced FA signaling compared to human fibroblasts, which may contribute to the phenotypes observed here of enhanced adhesive and motile properties in chimpanzee fibroblasts.

One of several of the major disparities in disease occurrence between humans and chimpanzees is that of incidence and progression of metastatic cancer. Though we clearly lack the same extensive population-level understanding of cancer incidence in chimpanzees, the data we do have from necropsies of captive chimpanzees indicates that epithelial cancer occurs at a rate of ≤ 4%, in sharp contrast to the > 20% of human deaths due to epithelial neoplasms, such as breast or lung carcinomas [5, 41-44]. Of further biomedical relevance, there is anecdotal evidence that humans and chimpanzees also differ in wound healing [45]. Intriguingly, these two disease states are linked – tumors have been described as wounds that do not heal [15] and gene expression profiles of cancerous tissue mimic that of tissue undergoing wound healing [46, 47]. The interspecies disparities in wound healing and cancer incidence may be due to alterations in the basic biological processes common to both of them, namely, cell adhesion and migration. Cancer is complex disease that results from aberrations of some of the most fundamental biological processes, including cell growth, proliferation, and migration. Fundamental differences in these processes between humans and chimpanzees may help explain to some extent the significant differences in cancer-related deaths. Several integrins expressed in epithelial cells influence metastasis, though whether increased or decreased expression in cancerous compared to normal tissue is inconsistent across integrins and cancer types [48]. Both of the integrins with increased expression in chimpanzee fibroblasts have been linked to metastasis in cancerous cell lines [49-52]. Reduced expression of ITGA7 was implicated with greater cell motility and metastasis [53] and thus there is potential that increased integrin expression in chimpanzee fibroblasts does contribute to decreased cell motility and potentially, decreased metastasis of cancerous cells. All of the cadherins up regulated in human (CDH2, CDH4) and chimpanzee (CDH1, CDH3) are implicated in cancers of various kinds (Roy 2014). CDH1 and CDH3 expression typically overlap (Roy 2014).

Furthermore, extracellular matrix proteins important for cell adhesion and migration include laminins and collagens. Collagen IV is the main collagen composing the basement membrane. COL4A6 was upregulated in chimpanzees at all timepoints (SI Table 2). However, during the serum challenge, humans upregulated COL6A1, COL6A3, and LAMA4 while chimpanzees upregulated COL9A3 in addition to their always upregulated COL4A6. This suggests that during a wound healing response, both species will upregulate secretion of basement membrane proteins of the ECM of similar families, but not the same genes. Finally, syndecans are transmembrane glycoproteins whose extracellular domains often act as co-receptors [54]. Interestingly, in normal fibroblasts, humans upregulated syndecan-3 (SDC3), a syndecan primarily found in adults in neuronal tissues (Table 1, SI Table 2) [54]. Chimpanzees upregulated SDC2 at all timepoints and SDC4 in normal fibroblasts, both of which are broadly expressed syndecans across multiple tissue and cell types [54]. This is in line with previous findings that altered gene expression in humans is enriched for neural processes [54]. This may represent a shared evolutionary pressure for increased neural activity.

## CONCLUSIONS AND IMPLICATIONS

The degree of change in coding sequence and protein function between humans and chimpanzees has long known to be insufficient to explain the stark phenotypic differences between these species [2, 4, 5]. Alterations in gene expression are known to influence human-specific phenotypes, including some that influence disease [1, 3, 4, 55]. However, this is one of few studies demonstrating that these divergences in expression can manifest in fundamental differences in cell biology. We demonstrate that there are functionally important differences at the cellular level between species corresponding that correspond with significant interspecies divergence in gene expression in wild-type unchallenged fibroblasts as well as those undergoing an *in vitro* mimic of both wound healing and cancer. Our study represents one of relatively few investigations into the basic cellular differences between species that may allow us to better understand evolutionary of complex disease traits

## FIGURES & TABLES

**SI Figure 1.**
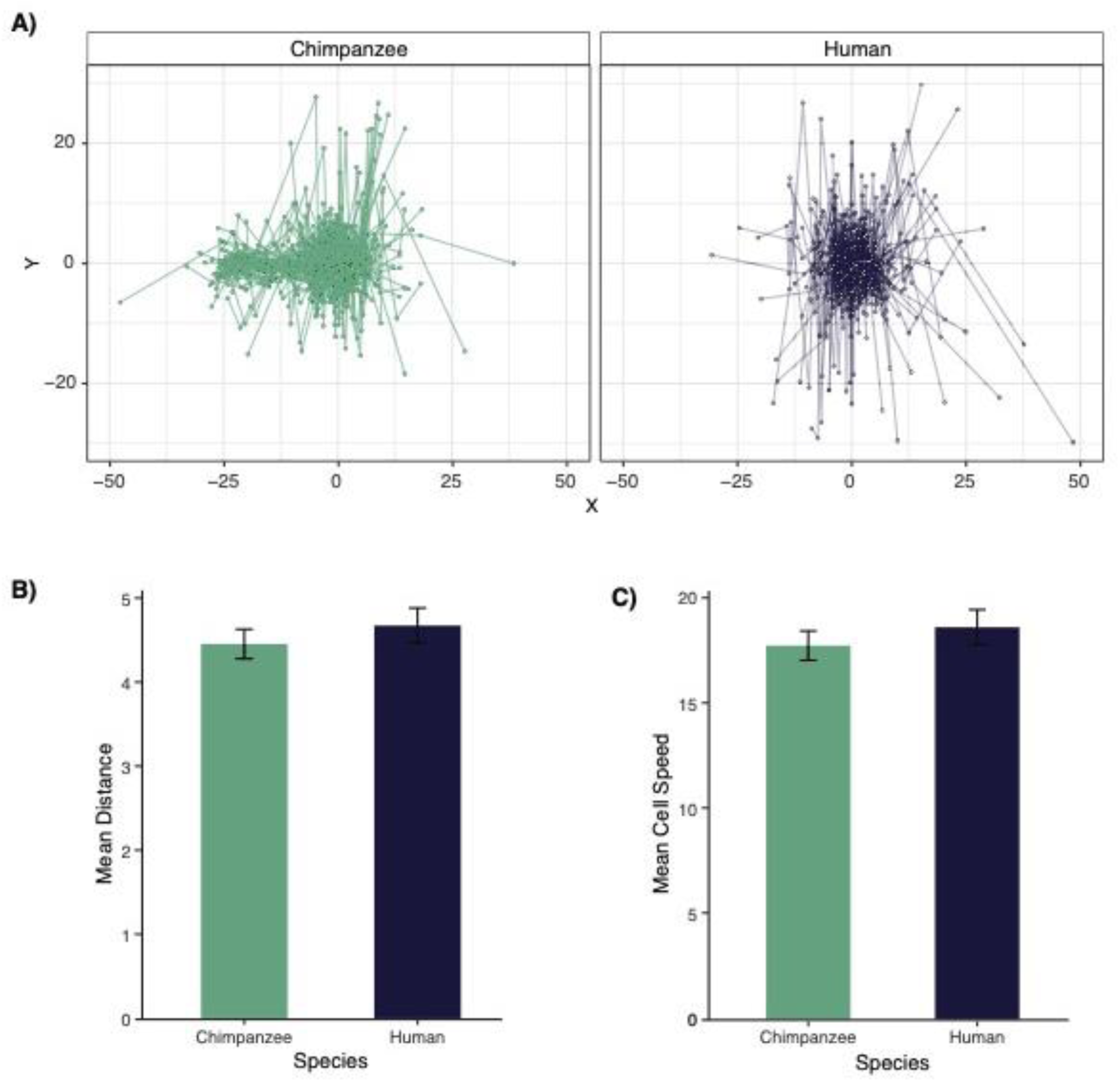
Human and chimpanzee fibroblast migration. Migration patterns of all observed human and chimpanzee fibroblasts plotted using X & Y coordinates obtained from FIJI (A). Species means for total distance traveled (B) and speed (C) of individual fibroblasts.

